# High-throughput phenotyping reveals multiple drought responses of wild and cultivated Phaseolinae beans

**DOI:** 10.1101/2024.02.09.579595

**Authors:** Jon Verheyen, Stijn Dhondt, Rafael Abbeloos, Joris Eeckhout, Steven Janssens, Frederik Leyns, Xavier Scheldeman, Veronique Storme, Filip Vandelook

## Abstract

Crop production worldwide is increasingly affected by drought stress. Although drought tolerance of a plant may be achieved through morphological, structural, physiological, cellular, and molecular adaptations, most studies remain limited to quantifying the effect of drought on biomass yield. Using a high-throughput phenotypic imaging system, we evaluated the drought tolerance of 151 bean accessions (Phaseolinae; Fabaceae) by quantifying five different traits simultaneously: biomass, water use efficiency, relative water content, chlorophyll content, and root/shoot ratio. Since crop wild relatives are important resources for breeding programmes, both wild and cultivated accessions were analyzed, the majority never evaluated for drought tolerance before. We demonstrate that the five traits are affected very differently by drought in the studied accessions, although a cluster analysis grouped the accessions into five distinct clusters with similar responses. We correlated the results for each accession to local climate variables at their original collection sites. Except for the root/shoot ratio, the results of all indicators were related to precipitation data, confirming that drought tolerant accessions grow in arid environments. This broader knowledge on the complex responses of plants to drought stress may prove an invaluable resource for future crop production.

**Highlight:** This study presents an innovative approach for the fast evaluation of different drought tolerance traits of legumes. Multiple responses to drought were observed in the economically important Phaseolinae beans.

## Introduction

Drought stress is one of the most important abiotic stresses affecting crop yield worldwide (Daryanto et al., 2016). Climate projections suggest that the Earth’s temperature will continue to rise throughout this century, leading to more frequent extreme climate events, including heatwaves, heavy precipitation, and drought (IPCC, 2021). Consequently, there is an expected increase in severe drought episodes, posing a serious threat to crop cultivation. Therefore, investigating the drought tolerance of cultivated food crops has become a paramount area of scientific research.

A plant’s response to drought stress is multifaceted and can be expressed in various ways. Two primary strategies to cope with drought are drought escape, characterised by a rapid and short life cycle, but an inability to withstand prolonged drought periods, and drought evasion, where the plant restricts water uptake or growth to avoid depleting soil moisture (Shantz, 1927). Coping with drought is achieved through morphological, physiological, molecular, and structural adaptations in plants. Although the capacity of a plant to resist drought is determined by all of these adaptations and the interactions between them, studies dealing with drought stress and plant cultivation focus mainly on the effect of drought on yield, which is considered the most valuable trait for food production. However, there are multiple other ways to quantify plant responses to drought, which can cover different aspects of the drought response spectrum.

Evaluating drought tolerance is often based on measurements of yield or biomass under drought stress compared to a control group. However, this approach presents challenges, as smaller plants may retain a higher biomass relative to their control group than larger plants (Iseki et al., 2014). One solution to this issue is to look at water use efficiency (WUE) instead of using absolute biomass or yield. WUE is the amount of biomass produced per unit of water used by the plant (Briggs and Shantz, 1913). Since lower growth rates are associated with a lower water consumption, WUE alleviates the bias towards smaller plants. In agricultural studies, WUE is therefore a more commonly used index (e.g. Anyia and Herzog, 2004). Drought stress also affects plants by limiting the amount of water available to different tissues. A decrease in relative water content (RWC) lowers the water potential and can increase the temperature in tissues, which in turn can lead to lower photosynthetic rates (Siddique et al., 2000). As the RWC of different plants may be affected very differently under drought stress, due to differences in water uptake and transpiration, RWC can also be used as a trait for drought tolerance screening (Schonfeld et al., 1988; Siddique et al., 2000; Belko et al., 2012). Several different methods to estimate RWC exist. The equation from Schonfeld et al. (1988) accurately calculates the RWC of plant tissues, although this method requires weighing the tissue before and after prolonged soaking and after oven drying, making it time-intensive and unsuitable for high-throughput purposes. Spectroscopic measurements on the other hand provide a more efficient approach. Water absorbs short-wave infrared light (900 nm – 2500 nm), and as such, the RWC of plant tissue can be correlated to the reflectance spectra in these wavelengths (Kim et al., 2015).

At the molecular level, one of the processes that is triggered by drought stress is an elevated production of H_2_O_2_, O_2-_ and other reactive oxygen species (de Carvalho, 2008). Within the chloroplasts, these lead to the degradation of chlorophyll, resulting in adverse effects (de Carvalho, 2008). A reduction in the relative chlorophyll content causes a lower photosynthetic rate (Fleischer, 1935) and has been associated with yield loss (Borrell et al., 2000). Through fluorescence measurements, Mathobo et al. (2017) demonstrated a decrease in chlorophyll content of *P. vulgaris* under different drought treatments. Vegetation mostly reflects electromagnetic waves in the near-infrared range (780 nm - 2500 nm), while chlorophyll strongly absorbs red light. As a consequence, spectroscopic indices making use of these wavelengths can be used to estimate the relative chlorophyll content. The normalized difference vegetation index (NDVI) was originally proposed to be used in remote sensing as a means to non-destructively estimate the amount of green biomass in satellite imagery (Rouse et al., 1974). This index, which is calculated as the difference between near-infrared reflectance and red reflectance, divided by their sum, has since been adopted as an accurate estimator for relative chlorophyll content in phenotyping studies (Bell et al., 2004) and as an indicator of drought stress (e.g. Trapp et al., 2016; Condorelli et al., 2018; Javornik et al. 2023).

Finally, it is important to also look at the root system to gain a more comprehensive understanding into the plants’ capacity to withstand drought conditions (Qi et al., 2019; Ma et al., 2021). Since an extensive, fast-growing root system can extract more moisture from the soil, plants with a large root system are often considered more drought tolerant. To eliminate a bias towards plants that are naturally large, the ratio between root biomass and shoot biomass is often used as an indicator. For many crops, including *P. vulgaris*, it has been shown that investing more energy into development of the root system relative to the shoot, results in a higher drought tolerance compared to plants with a low root/shoot ratio (Haider et al., 2012; Sofi et al., 2018). Seedlings of *P. vulgaris* experiencing drought stress have also been shown to alter their development towards a higher root/shoot ratio (Sofi et al., 2018). As such, this ratio can be used as a valuable predictor for drought resistance.

When economic, social, and infrastructural circumstances are considered, developing countries located in Africa, Asia, and Latin America are deemed most vulnerable to future drought disasters (Carrão et al., 2016). A critical component of these nations’ food security relies on the production and consumption of beans from the Phaseolinae subtribe (Fabaceae family), including the economically important *Vigna* and *Phaseolus* genera. These legumes rank prominently in agricultural land allocation for food crops. They account for the second largest share in Central America, followed by a third place in Africa and a fifth place in South America and Asia (FAO, 2020). The large majority of cultivated species within the Phaseolinae clade are restricted to the two genera *Vigna* and *Phaesolus*. The *Vigna* genus encompasses approximately 118 extant species, while *Phaseolus* comprises around 97 described species (WFO, 2022). The most well-known food crops include the common bean (*P. vulgaris*), cowpea (*V. unguiculata*), runner bean (*P. coccineus*), Bambara groundnut (*V. subterranea*), mung bean (*V. radiata*), lima bean (*P. lunatus*), tepary bean (*P. acutifolius*), adzuki bean (*V. angularis*) and the black gram (*V. mungo*).

Most of the species currently cultivated are found in moderately to extremely wet climates, but several crop wild relatives from these genera are located in arid environments where the annual precipitation is less than 400 mm (Fick and Hijmans, 2017). As the general awareness has considerably grown that crop wild genetic resources are of great potential in breeding programs, (Dempewolf et al., 2014), the current study characterises the degree of drought tolerance for several wild and cultivated bean species, aiming to provide a novel tool to increase drought tolerance in future bean cultivation. Apart from the study of Iseki et al. (2018), in which 69 *Vigna* accessions (including nine different cultivated species and 28 wild species) were screened, the degree of drought tolerance of many wild bean species has not been previously investigated. The results of Iseki et al. (2018) indicated that several wild accessions were highly drought tolerant, including accessions belonging to *V. subramaniana, V. trilobata, V. vexillata* var. *ovata* and *V. aridicola*. They also demonstrated that different genotypes from the same species can exhibit highly variable levels of tolerance (Iseki et al., 2018). For *Phaseolus*, no comparable large-scale studies have been performed to date.

The main aim of this study was to expose different responses to drought stress in a wide diversity of wild and cultivated Phaseolinae beans, using five different drought tolerance indicators derived from measurements of a high throughput phenotyping platform. The various plant traits upon which the tolerance indicators in this study are based (i.e. biomass penalty, water use efficiency, relative water content, relative chlorophyll content, and root/shoot ratio), are all driven by different processes and are therefore expected to cover different areas of the drought response spectrum. In addition, given that wild plants are adapted to local climate conditions, we tested whether the different drought stress indicators are related to local climate conditions the accessions originally grew. Finding close correlations between the drought tolerance indicators and climate conditions at the site of origin, would validate the approach of using high-throughput phenotyping in controlled conditions for large-scale drought tolerance screening of bean crop wild relatives. We specifically addressed the following questions: (I) how do different indicator variables relate to each other?; (II) how much variation exists within and among subgenera?; (III) are there groups of accessions with similar drought responses?; (IV) do the drought tolerance indicators correlate with the climate at the site of origin?

## Materials and methods

### Phaseolinae accessions

The 151 Phaseolinae accessions studied, included 8 genera (all of which were formerly considered part of either *Vigna* or *Phaseolus*), 12 subgenera and 65 species (Table S1). All taxon names were checked against the World Flora Online database (WFO 2022 a,b). Classification in subgenera followed Maréchal et al. (1978), and was updated with information from the WFO database (WFO 2022 a,b). There were 23 accessions that are defined as cultivated and 128 accessions as wild. Seeds of each accession were obtained from the Meise Botanic Garden collection, Belgium. The accessions were selected in such a way that they covered a wide taxonomic and ecological diversity, with a focus on *Vigna* species (including former *Vigna* genera), but supplemented with *Phaseolus* species. The accessions covered a large variety of local precipitation conditions, ranging from arid (< 200 mm annual precipitation) to extremely wet (> 2500 mm annual precipitation). Seeds had been stored in long term storage conditions (-20°C and 15% relative humidity) for variable amounts of time, but all had high viability (> 80%) and vigour.

Climate data based on coordinates at collecting sites were retrieved from WorldClim using DIVA-GIS (Hijmans et al., 2001). WorldClim contains high-resolution climate data from the years 1970-2000, during which period the majority of our accessions were collected (Fick and Hijmans, 2017). WorldClim provides data on 19 different climate variables, many of which are highly correlated. Therefore, we selected five variables (isothermality [quantifies how large the day-to-night temperatures oscillate relative to the summer-to-winter (annual) oscillations], precipitation seasonality, temperature seasonality, annual mean temperature and annual precipitation), that were both intuitive and informative. Coordinates were derived from information on collecting locality, whenever such collecting locality was sufficiently precise to derive coordinates. As such, climate data were derived for 118 ‘wild’ accessions (Table S1).

### Drought treatment and soil moisture measurement

Ten seeds per accession were scarified with a scalpel to break seed coat dormancy. Each seed was sown in a separate transparent plastic P12 pot and arranged in the greenhouse following a completely randomised design. The potting soil was supplemented with 3 g/L Osmocote® fertiliser. All pots were watered on flooding tables every 2-3 days to maintain a constant soil moisture at field capacity until 18 days after sowing (DAS), when the drought treatment started. On this date, the drought treatment started for five out of the ten pots from each accession, while the other five pots served as a control group (schematic representation in Fig. 1). The pots in the control group were watered every 1-2 days to maintain field capacity. The drought treatment involved a complete cessation of watering, resulting in a progressively increasing drought stress over time. Soil moisture was measured every 3-4 days until the pots were almost completely dry, as indicated by a threshold value (lower than 20%) of the ACO MMS-0 contactless moisture sensor based on high-frequency dielectric shift.

**Fig. 1.**
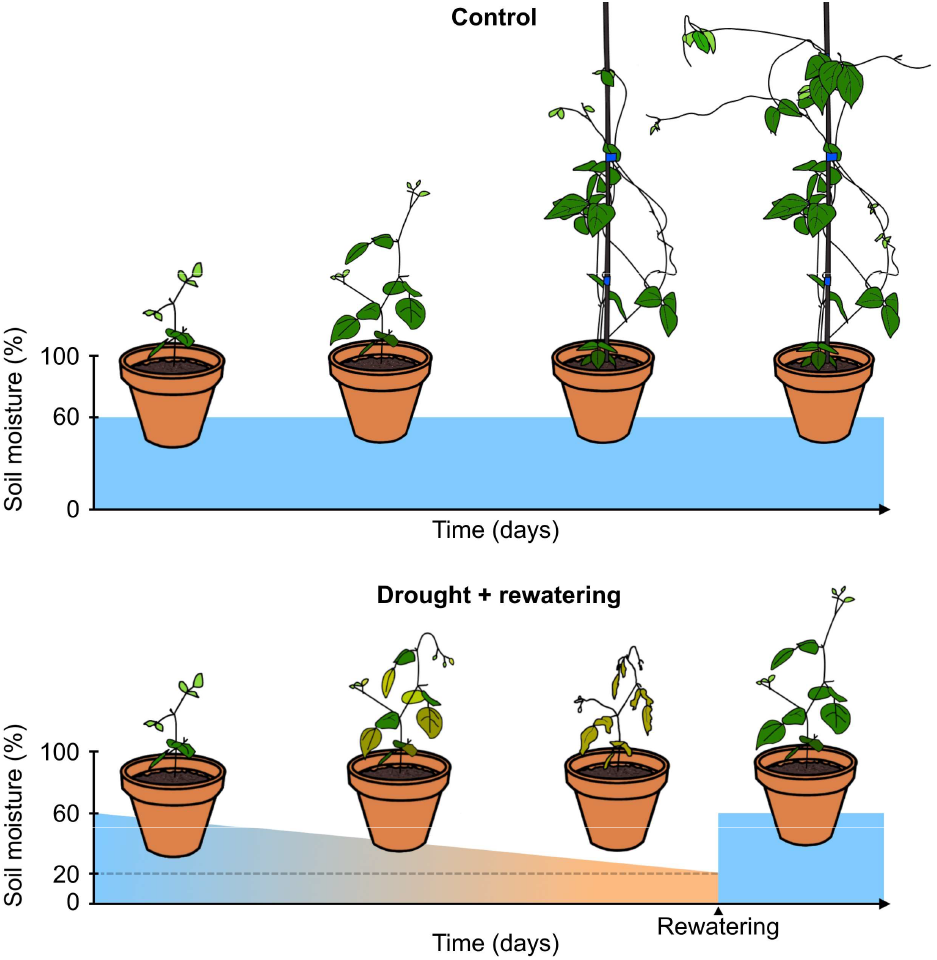
Watering scheme for drought treatment pots and control pots. Control pots were kept well-watered for a relatively constant soil moisture value of 60%. Pots in drought treatment were not watered until they reached a soil moisture value lower than 20%, at which point they were rewatered.

When this threshold was reached, pots were rehydrated until a soil moisture around field capacity was re-established, to allow for recovery of the plants. At 43 DAS, the final pots had dried out to the threshold value, were rewatered, and were allowed to recover for 3-4 more days, before the final measurements on 46 DAS. A day/night cycle with 13 hours of light was provided by SON-T lamps with an approximate photosynthetic photon flux density of 250 μmol/m2s photosynthetically active radiation. During the growth of the plants, day temperature was kept at a minimum of 20 °C and night temperature at 18 °C. A relative humidity of approximately 65 % was maintained at all times.

### Phenotypic imaging

Every 3 to 4 days starting from 18 DAS, we passed all plants through a high-throughput imaging system located at the VIB Agro-Incubator. Imaging of the drought-treated plants lasted for the entire length of the drought treatment, and one final imaging session was performed for each drought plant after rehydration. Since plants dried out at different rates, the amount of imaging sessions varied per pot. Imaging of the well-watered accessions lasted for a similar timespan as their corresponding drought-treated accessions. After being placed on conveyor belts, each plant passed by RGB, multispectral and hyperspectral cameras as well as a soil moisture measuring unit. Shoot biomass estimates were determined as the average projected area in side-view of the shoot on six images, with a 27° axial rotation of the plant between each image. The projected shoot area in side-view was photographed by a Prosilica GT 4905C RGB camera. The linear relationship between the projected side area and the aboveground biomass (R2 = 0.83, Fig. S1) was determined in a preliminary experiment based on 20 accessions with 8 to 10 plants for each accession using the *reg* procedure in SAS Studio (version 5.2, SAS Institute Inc., 2019).

An identical camera in bottom-view was used to determine the projected area of the roots on the bottom of the transparent pots as a proxy for root biomass. The reflectance of the plants at the wavelengths 1417 nm and 1528 nm for the estimation of RWC was determined by a Specim SWIR-CL-400-N25E hyperspectral camera (1000-2500 nm, with a spectral resolution of 3 nm) in top view (Kim et al., 2015). NDVI values were obtained from a multispectral FluxData FD-1665 3CCD camera containing two monochrome cameras for 680 nm and 800 nm. All experimental metadata, images, and phenotypic measurements were managed by the PIPPA software (version 0.14.20, VIB, 2021).

### Calculating drought tolerance indicators

For each bean accession, four drought tolerance indicators were determined based on aboveground biomass, WUE, RWC and NDVI. In addition, we calculated the root/shoot ratio which may be indicative of how well plants are adapted to drought. Biomass penalty was calculated as the proportion of aboveground biomass lost, due to the drought treatment compared to the control group ((biomass_control_ – biomass_drought_) / biomass_control_), as estimated by the projected area in side view and averaged over the plants belonging to the same accession and undergoing the same treatment (Fig. S2A). We used the biomass estimates of the drought plants after rewatering and the corresponding control plants, for the most accurate comparison, avoiding drought-induced leaf rolling. A low biomass penalty indicates a more drought tolerant accession.

For the indicator based on WUE, we followed the changes in WUE of the plants under drought stress. Since the pots did not receive any water during drought stress, we used the decrease in soil moisture as a proxy for the amount of water taken up by the plant. WUE values of the plants for each time interval, calculated as (biomass_t2_ - biomass_t1_) / (soil moisture_t1_ – soil moisture_t2_), were plotted against the mean soil moisture value over that time interval ((soil moisture_t1_ + soil moisture_t2_) / 2, further simply denoted as the soil moisture value). A positive WUE value for a chosen time interval indicates an increase in projected side-view area over that time interval and thus plant growth, while a negative value indicates a decrease and thus plant wilting. WUE values were plotted against soil moisture for each plant belonging to the same accession and a constrained generalised additive model was fitted for each accession using the *cgam* package in R (version 1.17, Liao and Meyer, 2019), assuming a smooth increasing curve with three knots and a gaussian distribution (Fig. S2B). The soil moisture value at which the plants ceased growing and started wilting could be found as the intercept of the fitted curve with the x-axis (WUE = 0). This x-intercept, the soil moisture at wilting, served as our WUE based indicator. A low value for this indicator implies that plants of the chosen accession only started wilting relatively late into the progressively worsening drought stress, thus suggesting a drought tolerant accession.

The third drought tolerance indicator, based on RWC, the soil moisture at leaf desiccation, was determined as the soil moisture value at which relative water content starts decreasing significantly. RWC values (reflectance_1528 nm_ / reflectance_1417 nm_) (Kim et al., 2015) were plotted against their corresponding soil moisture values and a linear plateau regression model was fitted to the data with the Levenberg-Marquardt algorithm from the *nlraa* package in R (version 1.9.3, Miguez, 2019) using the *SSlinp* self-starting function. The junction point between the two linear sections of the plateau regression indicated the soil moisture at leaf desiccation (Fig. S2C). A low value implies that the plants’ leaves started losing their water content relatively late into the drought stress, suggesting a drought tolerant accession.

A fourth indicator, based on NDVI, was equivalent to the indicator based on RWC, but was derived from the level of chlorosis presumed to be related to varying soil moisture levels. The NDVI values ((reflectance_680 nm_ – reflectance_800 nm_) / (reflectance_680 nm_ + reflectance_800 nm_)) were plotted against the corresponding soil moisture values (Fig. S2D) and a linear plateau regression model was fitted as described before. The soil moisture value at which NDVI started dropping significantly was determined as the junction point of a linear plateau regression. A low value for soil moisture at chlorosis implies a drought tolerant accession.

Finally, the root/shoot ratio was calculated as the average root/shoot ratio of all plants of the same accession following the same treatment on 18 DAS (Fig. S2E). We only used measurements before onset of the drought treatment, since measurements of the projected root area in pots under drought were unreliable. A high root/shoot ratio signifies plants that invest a relatively large amount of their growth into deeply penetrating roots relatively early in their development, implying a potentially greater capacity to resist drought stress.

### Statistical analyses

All analyses were performed with the R statistical software (version 4.2.2, R Core Team, 2022) in RStudio (version 2022.07.2 build 576, RStudio Team, 2022) or SAS Studio (version 5.2, SAS Institute Inc., 2019). Descriptive statistics, including mean, standard deviation, variance and range were calculated for all five drought tolerance indicators (n_accessions_ = 151). Spearman rank correlations were estimated between all indicator variables. These analyses were performed using the R base package.

To compare the proportion of within-subgenus variability and between-subgenus variability to the total variability, a mixed model analysis was performed on each indicator with subgenus as random effect. In addition, heterogeneous within-subject variability was modelled by estimating a different residual variance for each subgenus. This was done for each drought tolerance indicator separately. Subgenera with 3 or fewer accessions were omitted from this analysis. Mixed model analysis was performed with the mixed procedure from SAS/STAT in SAS Studio.

151 accessions with data for all 5 drought tolerance indicators were clustered with the Ward clustering method based on Manhattan distances using the *dist* and *hclust* function from the R *stats* package. The tree was subsequently cut in five clusters using the *cutree* function from the *dendextend* package (version 1.17.1, Galili, 2015). Visualisation of the tree was done with the *factoextra* (version 1.0.7, Kassambara and Mundt, 2020) package. Boxplots were generated by cluster membership.

A principal component analysis (PCA) was performed with 151 accessions to visualise the relationships between the different drought tolerance indicators using the *ggbiplot* (version 0.6.2, Vu, 2011), and *ggplot2* (version 3.4.2, Wickham, 2016) R packages. We also visualised the distribution of the different clusters obtained from the cluster analyses and wild versus cultivated accessions on the first two PC axes. Given that the data of the different indicator values covered similar ranges, no data transformations were performed prior to PCA analyses.

An RDA analysis was performed using the R *vegan* package (version 2.6-4, Oksanen et al., 2022) to visualise how climate conditions at the site of origin relate to the drought tolerance indicator variables. The analysis was carried out with 118 accessions for which climate data were available (Table S1). As mentioned before, five climate variables (isothermality, precipitation seasonality, temperature seasonality, annual mean temperature and annual precipitation) were included in the analyses together with five drought tolerance indicator variables. The full RDA model was statistically significant (P < 0.001, permutation test: #permutations = 999).

## Results

### Correlations between indicator variables

Mean values, ranges and standard deviation for the five drought tolerance indicators are listed in Table 1. The drought tolerance as expressed by WUE was significantly positively correlated to the biomass penalty (r = 0.39; P < 0.001) (Table 2, Fig. S3). RWC was significantly positively correlated with NDVI (r = 0.40; P < 0.001). All other correlations were not significant after Bonferroni correction (k = 10).

**Table 1.**
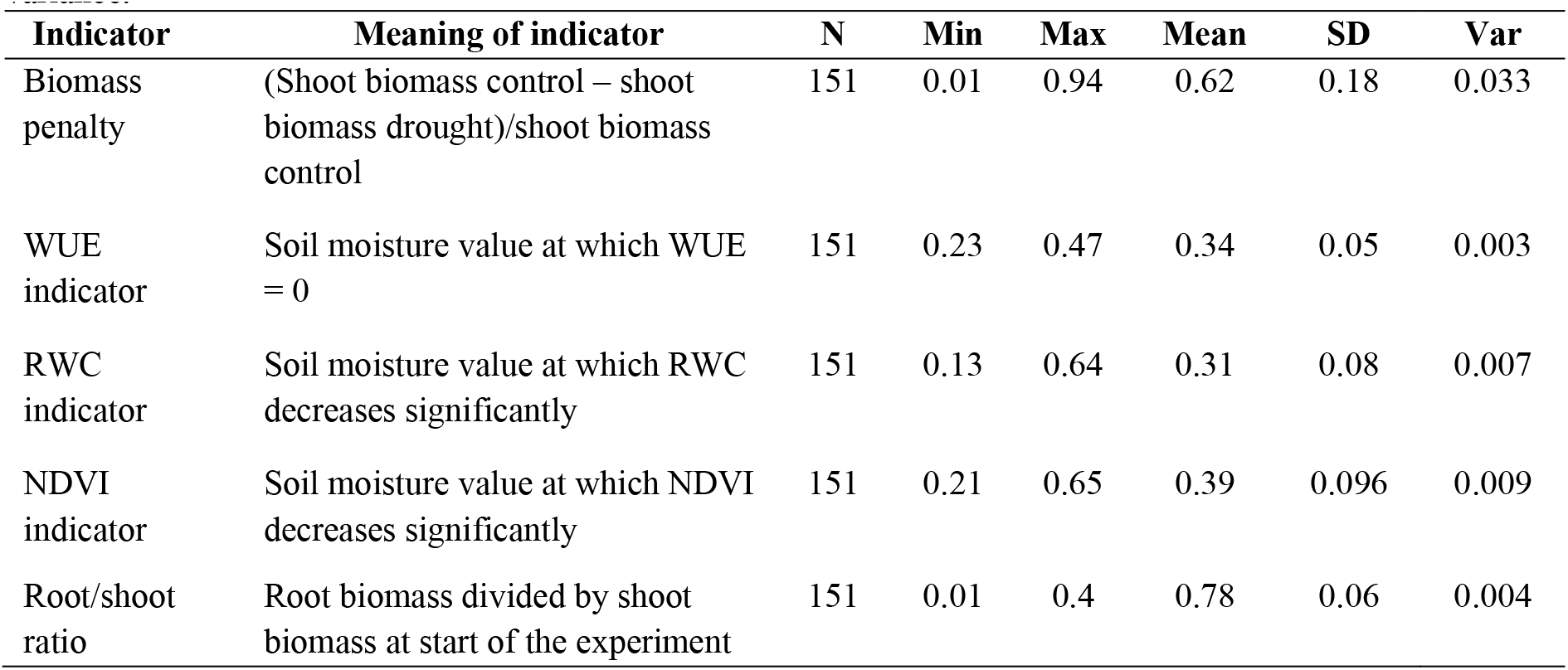
Descriptive statistics for all five moisture response indicators. N: number of accessions included, Min: minimum value, Max: maximum value, Mean: mean value across accessions, SD: ± 1 standard deviation, Var: variance.

| Indicator | Meaning of indicator | N | Min | Max | Mean | SD | Var |
| --- | --- | --- | --- | --- | --- | --- | --- |
| Biomass penalty | (Shoot biomass control – shoot biomass drought)/shoot biomass control | 151 | 0.01 | 0.94 | 0.62 | 0.18 | 0.033 |
| WUE indicator | Soil moisture value at which WUE = 0 | 151 | 0.23 | 0.47 | 0.34 | 0.05 | 0.003 |
| RWC indicator | Soil moisture value at which RWC decreases significantly | 151 | 0.13 | 0.64 | 0.31 | 0.08 | 0.007 |
| NDVI indicator | Soil moisture value at which NDVI decreases significantly | 151 | 0.21 | 0.65 | 0.39 | 0.096 | 0.009 |
| Root/shoot ratio | Root biomass divided by shoot biomass at start of the experiment | 151 | 0.01 | 0.4 | 0.78 | 0.06 | 0.004 |

**Table 2.** Spearman rank correlations (rho) between five moisture response variables across all accessions (n = 151). Asterisks indicate p-values (* < 0.05, ** < 0.001).

| Indicator | Biomass penalty | WUE indicator | RWC indicator | NDVI indicator | Root/shoot |
| --- | --- | --- | --- | --- | --- |
| Biomass penalty | 1 |  |  |  |  |
| WUE indicator | 0.40** | 1 |  |  |  |
| RWC indicator | 0.21* | 0.28** | 1 |  |  |
| NDVI indicator | -0.10 | 0.05 | 0.39** | 1 |  |
| Root/shoot ratio | -0.16 | -0.08 | -0.16* | -0.20* | 1 |

### Variation within and among subgenera

Within all subgenera except *Lasiospron*, most variation in the drought tolerance indicators was observed to be within-subgenus variation (Fig. 2; Table S2). Between subgenera, some clear differences were also observed (Fig. S4). The mean values of indicators based on WUE, RWC and NDVI were highest in *Ceratotropis* (0.38 ± 0.05; mean ± SD), *Sigmoidotropis* (0.48 ± 0.07) and *Condylostylis* (0.50 ± 0.10) respectively, while they were always lowest in *Plectotropis* (0.31 ± 0.05; 0.25 ± 0.07 and 0.33 ± 0.06, respectively). The mean indicator based on biomass penalty was highest in *Lasiospron* (0.82 ± 0.06) and lowest in *Sigmoidotropis* (0.41 ± 0.21). There was, however, a great amount of variation in biomass penalty within several subgenera (Table S2; Fig. S4A). The mean root/shoot ratio was highest in *Ceratotropis* (0.15 ± 0.12) and lowest in *Lasiospron* (0.04 ± 0.02). Especially for the indicators based on NDVI and biomass penalty, most variation was observed within subgenera (Table S2).

**Fig. 2.**
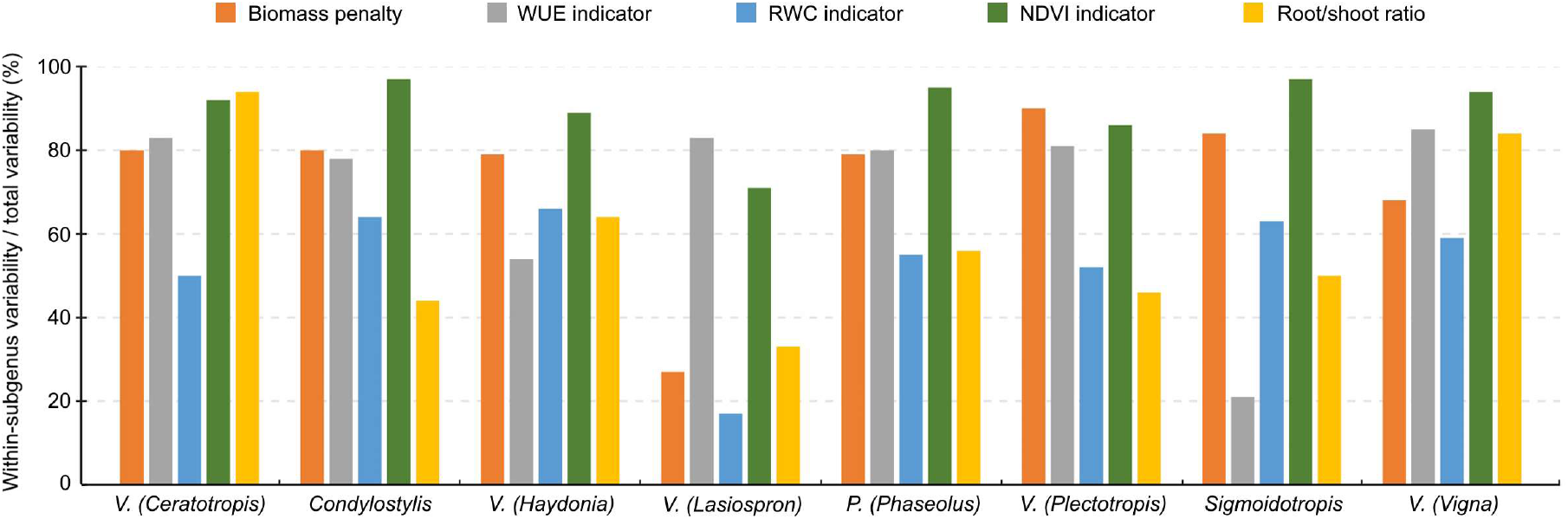
The proportion of within-subgenus variability/total variability for five drought response indicator values based on a mixed model analysis with subgenus as random effect. Heterogeneous within-subject variability was modelled by estimating a different residual variance for each subgenus. Subgenera with 3 or fewer accessions were omitted.

### Grouping bean accessions with similar drought responses

Cluster analyses enabled us to group accessions into five clusters of accessions with similar drought responses within clusters (Fig. 3). Clusters 1 and 2 (Fig. 3) were characterised by high values for biomass penalty (Fig. 4A), which may indicate that these are tall plants whose growth is considerably reduced by drought. Accessions in cluster 2 also showed a higher root/shoot ratio as compared to other accessions (Fig. 4E), indicating that their growth is affected by drought despite a higher root/shoot ratio. Accessions in cluster 3 had a high indicator value for RWC and NDVI (Fig. 3, Fig. 4C and 4D), which means that the response was mainly visible in the form of reduced photosynthesis. The accessions in cluster 4 had low values for the WUE and RWC indicator (Fig. 3, Fig. 4B and 4C), meaning they responded better to drought as compared to accessions in other clusters. Finally, accessions in cluster 5 had high values for the WUE indicator (Fig. 3, Fig. 4B) and low values for all other indicator variables.

**Fig. 3.**
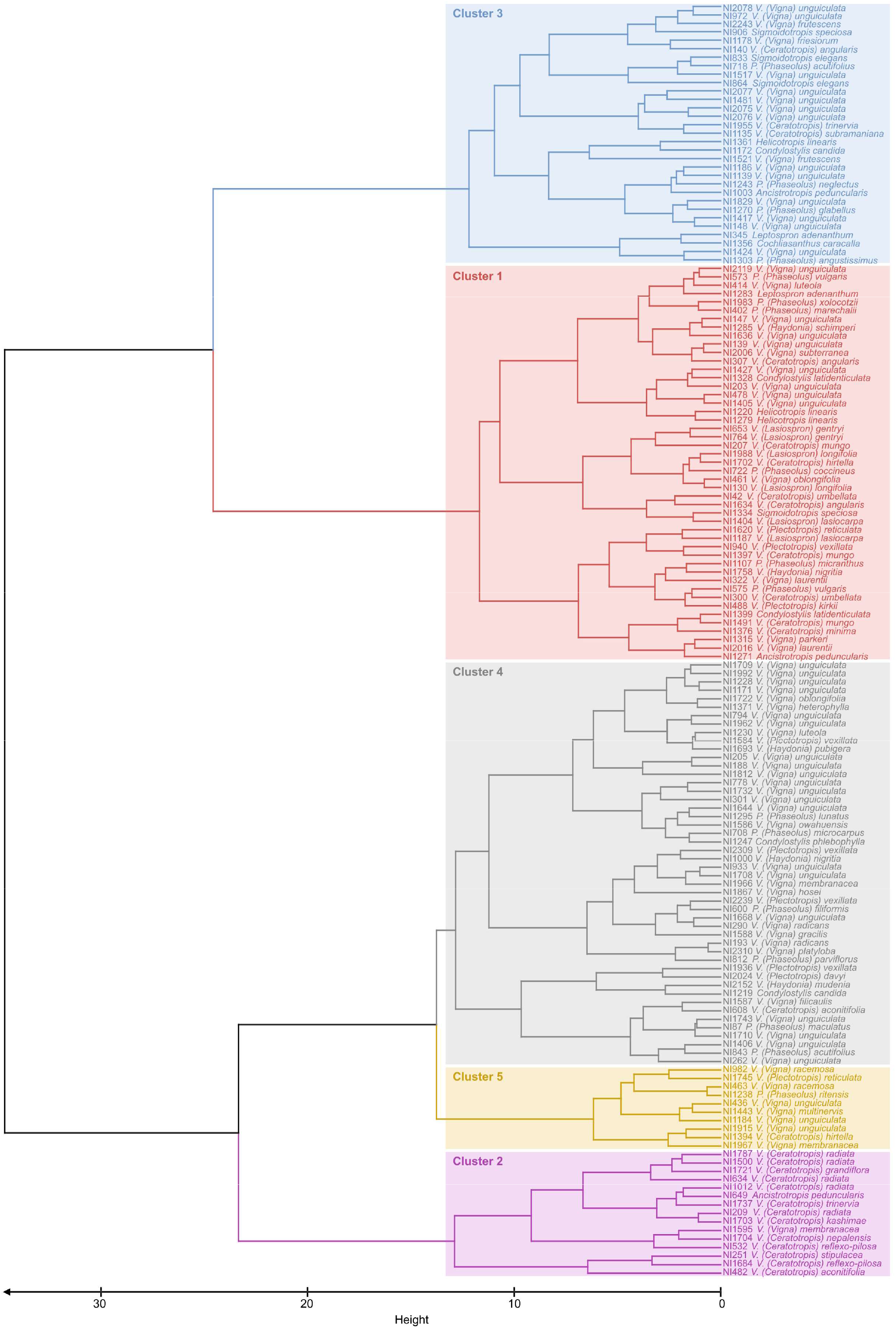
Dendrogram obtained by clustering analyses on 151 bean accessions with data for five drought response indicators. The Ward clustering method was used based on Manhattan distances. The tree was cut in five clusters, grouping accessions with similar drought responses.

**Fig. 4.**
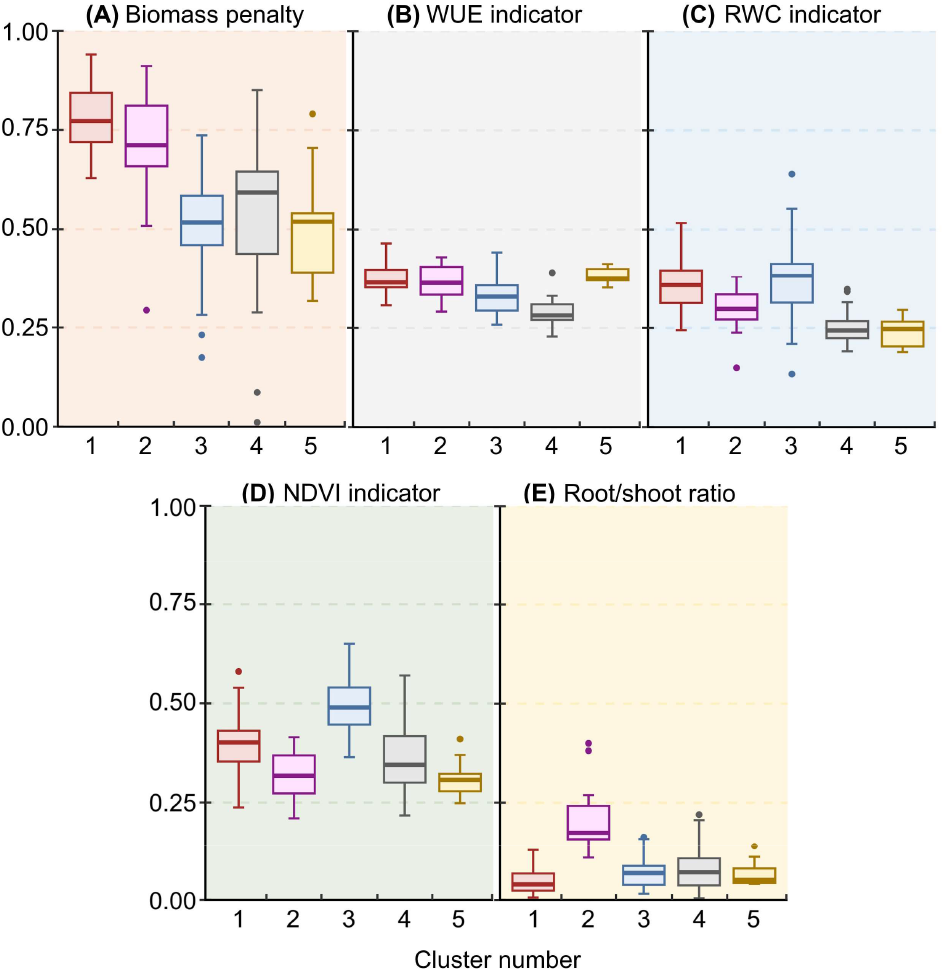
Boxplots of median drought response indicator values for each of five clusters generated by a cluster analysis on 151 bean accessions.

A total of 151 bean accessions were included in the PCA (Fig. 5). The first three PCs explained 33.5, 26.8 and 16.9% of the variance. The PCA confirmed the positive correlation between the WUE indicator and the biomass penalty and between RWC and NDVI indicators. It further suggests a negative correlation between the root/shoot ratio and the NDVI indicator and between the root/shoot ratio and RWC indicator despite the negligible negative correlation found with the Spearman rank correlation (Table 1). WUE indicator and biomass penalty were relatively independent from root/shoot ratio and NDVI indicator, hence they were likely situated in other areas of the drought tolerance spectrum. Generally, accessions situated at the right hand side of PC1 could be considered more drought tolerant, as a high value for the indicators based on NDVI, RWC, WUE and biomass penalty were all indicative of a poor tolerance to drought. The different clusters derived from the cluster analyses separated very well along PC1 and PC2. Overall, accessions in cluster 1 and 2 seemed to respond less well to drought, while accessions in cluster 3, 4 and 5 are more drought tolerant. Nonetheless, considerable variation in drought tolerance was observed within clusters. When cultivated and wild accessions were coloured separately in a PCA (Fig. S5), no obvious differences in drought tolerance between wild and cultivated accessions could be observed. The cultivated accessions were distributed somewhat continuously along the entire PC1 axis.

**Fig. 5.**
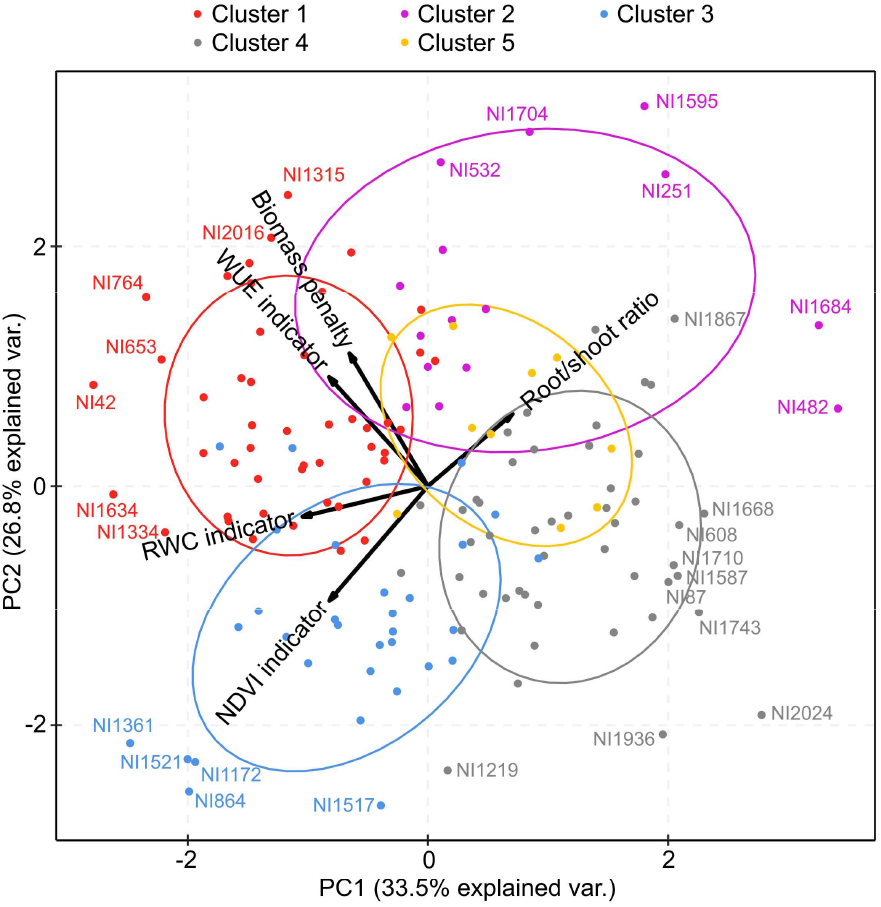
Two first axes of a principal component analysis (PCA) performed with 151 accessions to visualise the relationships between the five different drought response indicators. The distribution of the five different clusters obtained from a cluster analyses based on drought response indicators is indicated with different color schemes.

### Relationship of drought tolerance with local climate

Only the first two axes of the RDA model were statistically significant (P < 0.001) and were retained (Fig. 6). The included climatic variables (isothermality, precipitation seasonality, temperature seasonality, annual mean temperature and annual precipitation) explained 14.2% (adjusted R2) of the variation in drought tolerance among genotypes. Only three of the climatic variables (isothermality, annual mean temperature and annual precipitation) were statistically significant (P < 0.001; P = 0.004; P < 0.001, respectively). Annual precipitation at the site of origin was significantly positively related to the indicators based on biomass penalty, RWC and WUE (Table 3). Annual mean temperature was significantly positively related to the root/shoot ratio, while temperature seasonality decreased significantly with the indicator based on biomass penalty. Finally, a significant positive relation was observed between isothermality and biomass penalty, while a significant negative relation with root/shoot ratio was observed (Table 3).

**Table 3.**
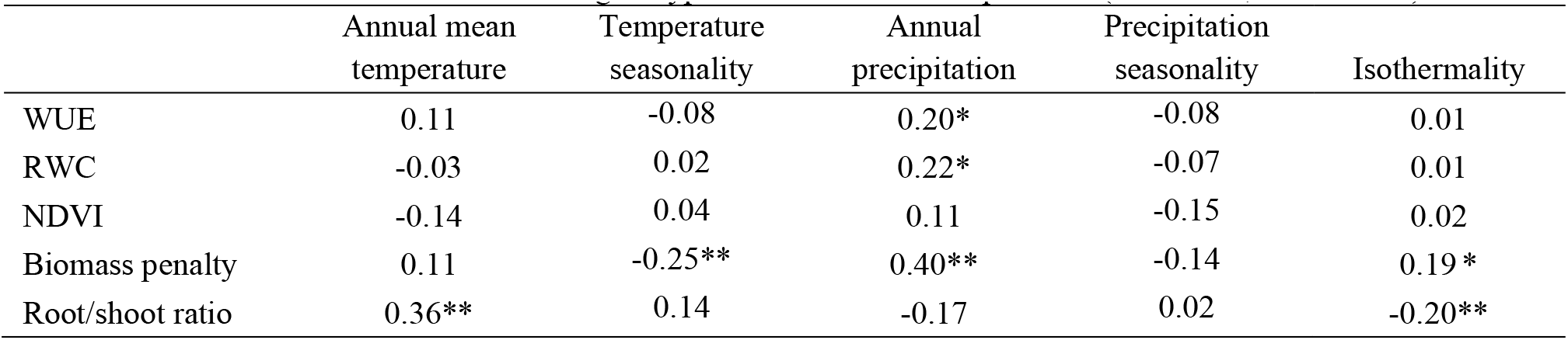
Pearson correlation coefficients for correlations between five drought response indicators and five climatic variables for 118 wild Phaseolinae genotypes. Asterisks indicate p-values (* P < 0.05; ** P < 0.01).

**Fig. 6.**
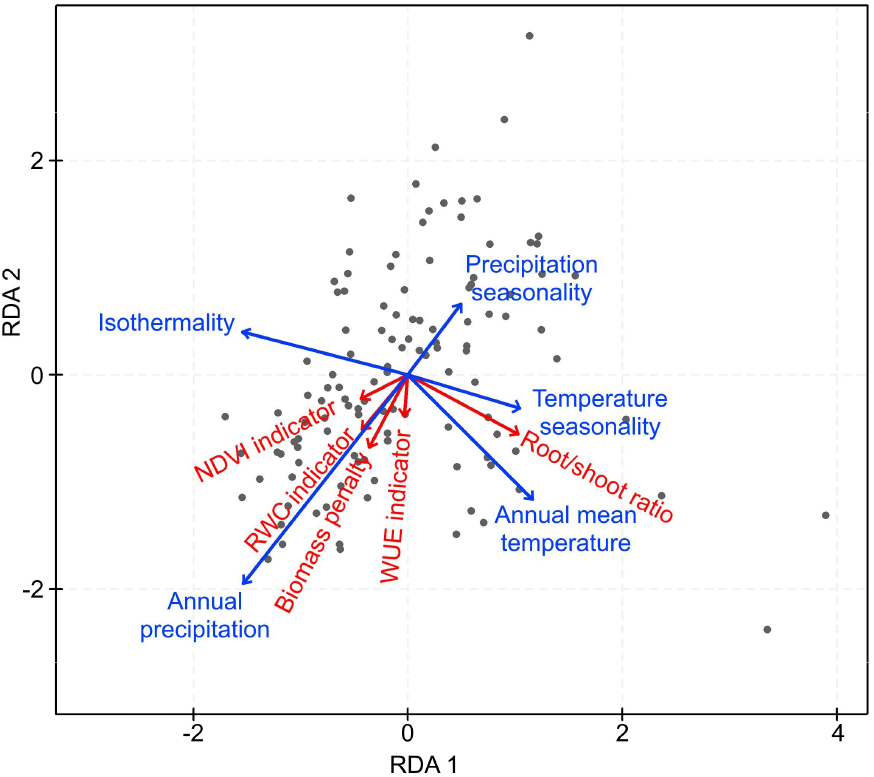
An RDA analysis visualising how climate conditions at the site of origin relate to the drought response indicator variables. The analysis was carried out with 118 accessions. Five climate variables (isothermality, precipitation seasonality, temperature seasonality, annual mean temperature and annual precipitation) were included in the analyses together with five drought response indicator variables.

## Discussion

By using five drought tolerance indicators to analyse the drought response of 151 Phaseolinae bean accessions, we were able to get a much more complete picture of drought tolerance across this agronomically important clade. Most of the indicators were not correlated significantly to each other, confirming that their underlying molecular and cellular mechanisms can respond very differently to drought stress. The RWC and NDVI indicators were moderately correlated, as well as the biomass penalty and WUE indicator, which was to be expected as these indicators were derived from similar measurements in both cases. Our ability to differentiate various drought tolerance clusters and the fact that the drought tolerance of wild Phaseolinae bean accessions did relate to the climate at the site of collection, indicates that a high-throughput phenotyping platform enables a cost-efficient screening of drought tolerances of highly diverse Phaseolinae genotypes as further discussed below.

The aim of studying multiple different indicators, on top of the biomass measurements that most drought tolerance studies are limited to, was to gain a broader understanding of the effect of drought stress on plant morphology, physiology and molecular processes. Overall, the underlying cellular and molecular processes are all negatively affected by drought stress, which is demonstrated by the decreasing trend in biomass, WUE, RWC and NDVI seen in virtually all accessions. However, subtle differences between the indicators were detected. Groups of accessions that show similar responses to drought based on the different drought response indicators, could also be discerned. There was, for example, one group of accessions (cluster 5) where the drought response was mostly manifested through WUE, while all other drought tolerance indicators revealed only a minor drought response. It has been argued that WUE might not provide much information about the competitive or yield advantage of one particular species over another because improved WUE may actually restrict growth (Hubick et al., 1986). The trait has, however, been studied quite often because it can give an idea of the variation amongst genotypes in ability where water is limiting (Anyia and Herzog, 2004). Some of the drought tolerance indicators are more similar to others, such as NDVI and RWC, which are closely related to the photosynthesis process and both derived from infrared reflectance measurements in our study. A strong correlation between NDVI and RWC has been observed in previous studies (e.g. Jiang et al., 2009). In our study, accessions in cluster 3 were characterised by a high indicator value for both NDVI and RWC, which means that one of these two drought tolerance indicators is probably redundant.

Root/shoot ratio was the drought tolerance indicator that deviated most from all other indicators. Root area measurements with the bottom-view camera proved inaccurate during drought stress, as the dried up soil formed clumps in the pots, pulling the roots away from the bottom of the pots. For this reason, we measured the root/shoot ratio before application of the drought stress, and it is therefore not a physiological response to drought stress but rather an indicator of potential drought tolerance. Out of the five indicators, root/shoot ratio provides the lowest contribution to the first two principal components in our PCA analysis. The fact that the root/shoot ratio is also the indicator that is the least related to local precipitation data, and seems to be more strongly related to annual temperature, suggests that the root/shoot ratio as measured here was not a sufficient indicator for drought tolerance. Further studies would benefit from methods to accurately estimate root biomass non-destructively during the drought stress period, as the root/shoot ratio may alter differently for different species under drought stress (Zhou et al., 2018). In the past, electrical capacitance has been used for non-destructive root measuring, and although this method is reliable in hydroponics and sand-based growth medium, measurements in soil-based experiments proved inaccurate (Aulen and Shipley, 2012). For now, non-destructive measuring of root biomass in soil-based experiments remains limited to imaging techniques, which are generally less accurate as only a small part of the root system can be imaged.

While many studies have focused on screening for drought tolerance in domesticated *Vigna* (e.g. Anyia and Herzog, 2004; Belko et al., 2012; Tsoata et al., 2015) and *Phaseolus* species (Souza et al., 2003; Rosales et al., 2012; Mathobo et al., 2017), far fewer studies have screened drought responses in the closely related wild relatives (but see Iseki et al., 2018). Cultivated species within the *Phaseolus* and *Vigna* genera face several environmental challenges that affect their global production, such as drought, extreme temperatures, and salinity (Beebe et al., 2012; Harouna et al., 2018). This is where wild relatives of crop plants may fill an important gap, as they often have a greater adaptability to abiotic stresses and are expected to have valuable genes for breeding (Acosta-Gallegos et al., 2007; Palmgren et al., 2014).

As far as we know, this is also the first study screening drought tolerance across different genera and subgenera within the agronomically important Phaseolinae clade. One potential pitfall of screening drought tolerance over such a wide diversity, is that different clades may have evolved different strategies to cope with drought stress. Although differences between subgenera were indeed observed, most variation was observed within subgenera. Even within species, considerable variation in drought tolerance was observed. Accessions of *V. unguiculata*, for example, were found across four different drought tolerance clusters, which confirms the findings of Iseki et al. (2018) for this species. On the other hand, accessions of *V. radiata* were remarkably similar in their drought response as all five of them clustered together in cluster 2. A large intraspecific diversity in drought tolerance makes comparisons between different studies not using the same accessions difficult, and shows the importance of using stable identifiers to characterise accessions and the exchange of information between seed banks across the world preserving and studying this material (Debouck, 2014; Nair et al., 2023). A few overall highly drought tolerant wild accessions worthy of mentioning are: *P. novoleonensis* (NI 1237), *V. vexillata* var. *lobatifolia* (NI 546), *V. kokii* (NI 2178), *V. reflexo-pilosa* var. *reflexo-pilosa* (NI 1684), *V. unguiculata* subsp. *stenophylla* (NI 1904), *V. unguiculata* subsp. *unguiculata* var. *spontanea* (NI 1668), and *V. davyi* (NI 2024). Of these, two were phenotyped by Iseki et al. (2018) as well. A different accession of *V. vexillata* var. *lobatifolia* was found to be moderately drought tolerant, while a different accession of *V. reflexo-pilosa* var. *reflexo-pilosa* was found to be relatively drought sensitive. This again reinforces the idea of significant intraspecific diversity.

The redundancy analysis showed that all drought tolerance indicators were to a varying extent related to climate at the original sampling locations of the wild accessions. A strong relationship was found between annual precipitation and the biomass-related indicators, biomass penalty, soil moisture at wilting (WUE) and with soil moisture at leaf desiccation (RWC). Root/shoot ratio was related to annual mean temperature at the site of origin. These results validate the use of the high-throughput phenotyping platform for screening drought tolerance across a wide diversity of Phaseolinae species and hint at stronger adaptation to drought for accessions and species that grow in dry regions. However, numerous other factors, such as vegetation structure, hydrogeography, altitude, and rhizobial and mycorrhizal interactions may influence the drought tolerance of legumes (Cortés et al., 2013). In future studies, the influence of other factors could be explored. Mainly rhizobial and arbuscular mycorrhizal interactions, which have been shown to significantly alleviate drought stress effects in Fabaceae, should be considered (Rasaei et al., 2013; Tsoata et al., 2015).

No clear distinction could be found in drought tolerance responses between cultivated and wild accessions. A PCA showed no separate grouping for wild and cultivated accessions. Moreover, cultivated accessions were found on both extreme ends of PC1 and in all five drought tolerance clusters. This indicates that within cultivated Phaseolinae, considerable variation exists in drought tolerance. This contrasts somewhat with the observations of Cortés et al. (2013), who found that wild common bean (*Phaseolus vulgaris*) occupies more geographical regions with extensive drought stress than the cultivated accessions. Iseki et al. (2018) also observed that the most drought tolerant wild *Vigna* accessions performed better than the domesticated accessions that were cultivated in drought-prone areas. For subsequent studies, a more balanced number of accessions, both wild and cultivated, should be phenotyped for the important crop species, such as the common bean (*P. vulgaris*), Bambara groundnut (*V. subterranea*), and the mung bean (*V. radiata*). For the predicted climate changes, these accessions could be crucial in maintaining crop production and increased drought is only one of the factors that hampers contemporary and future crop production.

High-throughput phenotyping platforms are increasingly used in agricultural settings as they come with several advantages. The main advantage is that large numbers of genotypes can be screened for abiotic stress responses, such as drought, in a time- and cost-efficient manner. We managed to screen the drought response for over 1500 plants from 151 accessions, with measurements taking place every 3 to 4 days. This would not have been possible with manual measurements. In addition, because the phenotyping platform is semi-automated, all measurements are relatively standardised and measurement error due to different observers was reduced. Using different types of cameras also allowed us to collect a huge amount of data variables, of which only a subset was used in the present analysis. With this study, we have shown that high-throughput phenotyping can also aid screening of wild genotypes, which are generally much more variable in terms of plant size and growth habit (Popoola et al., 2015) as compared to varieties within a single species. Within-subgenus and between-subgenus variation in drought tolerance was large in our study, but nonetheless groups of accessions with similar drought tolerances could be delimited.

There were also some limitations to the high-throughput phenotyping platform. The drought experiment was run for a limited amount of time, as plants grew too large, reducing measurement accuracy at some point. As many of the Phaseolinae accessions are climbers, they had to be tied up to sticks regularly. The type of stress provided was also quite severe and limited in time, which may not be representative of all natural situations. Plant response to a given water deficit is for example strongly dependent on the previous occurrence and intensity of other drought stress events and the presence of other stresses (Seleiman et al., 2021). A logical next step would be to run experiments with different types of water deficit treatments, such as for example a prolonged exposure to drought, and in interaction with variation of other environmental conditions, such as variation in temperature. In addition, tying the information obtained through our high-throughput phenotyping experiment with anatomical measurements (e.g. stomatal density and size), phenology (e.g. flowering and pod production), and functional traits (stomatal conductance and transpiration rate) related to stress tolerance will help to elucidate the role of each component in the adaptation to fluctuations in water deficit (Correia et al., 2022).

In conclusion, our study provides an explorative approach to simultaneously characterise the changes in selected morphological, physiological and molecular plant traits under drought stress. The indicators soil moisture at wilting, soil moisture at leaf desiccation and soil moisture at chlorosis, together with a biomass penalty, provide a broad and nuanced vision on the complexities of a plant’s drought response. We derived these indicators for 151 accessions of both wild and cultivated *Vigna* and *Phaseolus* bean species. Our study thus revealed accessions that may be of interest in breeding programmes. We demonstrated the presence of large intraspecific differences in the capacity to resist drought stress, which could be exploited in breeding. Lastly, we compared drought tolerance of the accessions with local climate data of their collection sites and found strong relationships with precipitation data, confirming that the most drought tolerant Phaseolinae accessions mainly grow in arid climates. In order to maintain or increase our current global bean production in times of increasing drought disasters, further studies should expand upon this knowledge gained from phenotypic traits. Both genomic information and studies on the rhizobial and mycorrhizal interactions could provide invaluable insights.

## Abbreviations

DAS: days after sowing
NDVI: normalized difference vegetation index
PCA: principal component analysis
RDA: redundancy analysis
RWC: relative water content
WUE: water use efficiency

## Supplementary data

The following data are available in the supplementary files.

*Table S1*. List of all 151 accessions with taxon name, country and coordinates where seeds were originally collected, cultivated or wild origin, and other data used in the analyses.

*Table S2*. The proportion of within-subgenus variability and between-subgenus variability to the total variability in drought response indicator values.

*Fig. S1*. The linear relationship between the projected side area and the aboveground biomass.

*Fig. S2*. Explanatory graphs for each of the five drought response indicators.

*Fig. S3*. Scatterplots of the five drought tolerance indicators in relation to each other.

*Fig. S4*. Boxplots of median drought response indicator values for different subgenera of *Vigna* and *Phaseolus* beans.

*Fig. S5*. Principal component analysis (PCA) performed with 151 accessions to visualise the relationships between five different drought response indicators.

## Acknowledgements

We thank Sarah le Pajolec, Yannick Coeckelberghs and Gery Van den Troost (Meise Botanic Garden) for practical help during the experiment. Prof. Johan Robben (KU Leuven) for supervision and feedback on the statistical analyses.

## Author contributions

XS, SD, FV, and SJ: conceptualization; JV, VS, SD, RA, JE, and FL: data curation; JV, VS, FV, and XS: formal analysis; JV, XS, SD, RA, JE, and FL: methodology; FV, and XS: supervision; JV, VS, and FV: visualization; JV, XS, VS, and FV: writing - original draft; SD, RA, JE, FL, and SJ: writing - review & editing.

## Data availability

Data will be shared on request.

