## Supplementary Figures S1-S5 and Table S2 for "High-throughput phenotyping reveals multiple drought responses of wild and cultivated Phaseolinae beans"

### Supplementary data

*Table S1.* See supplementary Excel file.

*Table S2.* The proportion of within-subgenus variability and between-subgenus variability to the total variability in drought response indicator values based a mixed model analysis with subgenus as random effect. In addition, heterogeneous within-subject variability was modelled by estimating a different residual variance for each subgenus.

| (Sub)genus | Within variability | Total variability | Within variability (%) |
|---|---|---|---|
| WUE indicator values | | | |
| *V. (Ceratotropis)* | 0.0025 | 0.0030 | 83% |
| *Condylostylis* | 0.0018 | 0.0024 | 78% |
| *V. (Haydonia)* | 0.0006 | 0.0011 | 54% |
| *V. (Lasiospron)* | 0.0026 | 0.0031 | 83% |
| *P. (Phaseolus)* | 0.0020 | 0.0025 | 80% |
| *V. (Plectrotropis)* | 0.0023 | 0.0028 | 81% |
| *Sigmoidotropis* | 0.0001 | 0.0007 | 21% |
| *V. (Vigna)* | 0.0028 | 0.0033 | 85% |
| RWC indicator values | | | |
| *V. (Ceratotropis)* | 0.0042 | 0.0083 | 50% |
| *Condylostylis* | 0.0073 | 0.0114 | 64% |
| *V. (Haydonia)* | 0.0082 | 0.0123 | 66% |
| *V. (Lasiospron)* | 0.0009 | 0.0050 | 17% |
| *P. (Phaseolus)* | 0.0050 | 0.0091 | 55% |
| *V. (Plectrotropis)* | 0.0046 | 0.0087 | 52% |
| *Sigmoidotropis* | 0.0070 | 0.0111 | 63% |
| *V. (Vigna)* | 0.0061 | 0.0102 | 59% |
| NDVI indicator values | | | |
| *V. (Ceratotropis)* | 0.0063 | 0.0068 | 92% |
| *Condylostylis* | 0.0197 | 0.0203 | 97% |
| *V. (Haydonia)* | 0.0046 | 0.0052 | 89% |
| *V. (Lasiospron)* | 0.0014 | 0.0019 | 71% |
| *P. (Phaseolus)* | 0.0114 | 0.0119 | 95% |
| *V. (Plectrotropis)* | 0.0033 | 0.0039 | 86% |
| *Sigmoidotropis* | 0.0160 | 0.0165 | 97% |
| *V. (Vigna)* | 0.0084 | 0.0090 | 94% |
| Biomass indicator values | | | |
| *V. (Ceratotropis)* | 0.039 | 0.0491 | 80% |
| *Condylostylis* | 0.0396 | 0.0495 | 80% |
| *V. (Haydonia)* | 0.0373 | 0.0473 | 79% |
| *V. (Lasiospron)* | 0.0037 | 0.0137 | 27% |
| *P. (Phaseolus)* | 0.0371 | 0.0470 | 79% |
| *V. (Plectrotropis)* | 0.0933 | 0.1033 | 90% |
| *Sigmoidotropis* | 0.0533 | 0.0633 | 84% |
| *V. (Vigna)* | 0.0211 | 0.0311 | 68% |
| Root/shoot ratio | | | |
| *V. (Ceratotropis)* | 0.0142 | 0.0150 | 94% |
| *Condylostylis* | 0.0007 | 0.0017 | 44% |
| *V. (Haydonia)* | 0.0016 | 0.0026 | 64% |
| *V. (Lasiospron)* | 0.0005 | 0.0014 | 33% |
| *P. (Phaseolus)* | 0.0012 | 0.0021 | 56% |
| *V. (Plectrotropis)* | 0.0008 | 0.0017 | 46% |
| *Sigmoidotropis* | 0.0009 | 0.0019 | 50% |
| *V. (Vigna)* | 0.0048 | 0.0057 | 84% |

*Fig. S1.* The linear relationship between the projected front area and the aboveground biomass was determined in a preliminary experiment based on 20 accessions with 8 to 10 plants for each accession using the *reg* procedure in SAS Studio version 5.2 (SAS Institute Inc., Cary, NC, USA).

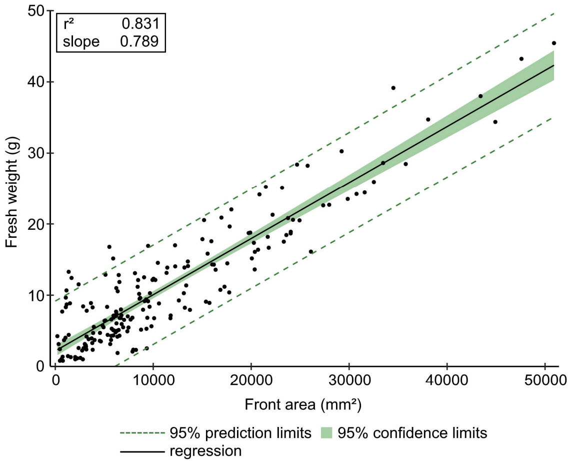


*Fig. S2*. Explanatory graphs for each of the five indicators. For each panel, the left figure gives an example of a drought tolerant accession, and the right figure gives an example of a drought sensitive accession, based on the indicator on that panel. (A) Projected side area of drought and control groups as an estimator for biomass. Biomass penalty was calculated as the difference of the means (indicated by crosses), divided by the mean of the control group. (B) WUE plotted against soil moisture and fitted with a CGAM model curve. Colors indicate individual plants. The intercept of the curve with the x-axis is the soil moisture at wilting. (C) RWC plotted against soil moisture and fitted with linear plateau regression. Colors indicate individual plants. The x-value of the junction point of the regression is the soil moisture at leaf desiccation. (D) NDVI plotted against soil moisture and fitted with linear plateau regression. Colors indicate individual plants. The x-value of the junction point of the regression is the soil moisture at leaf desiccation. (E) Root/shoot ratio. A cross indicates the mean, which is used as the fifth indicator for each accession.

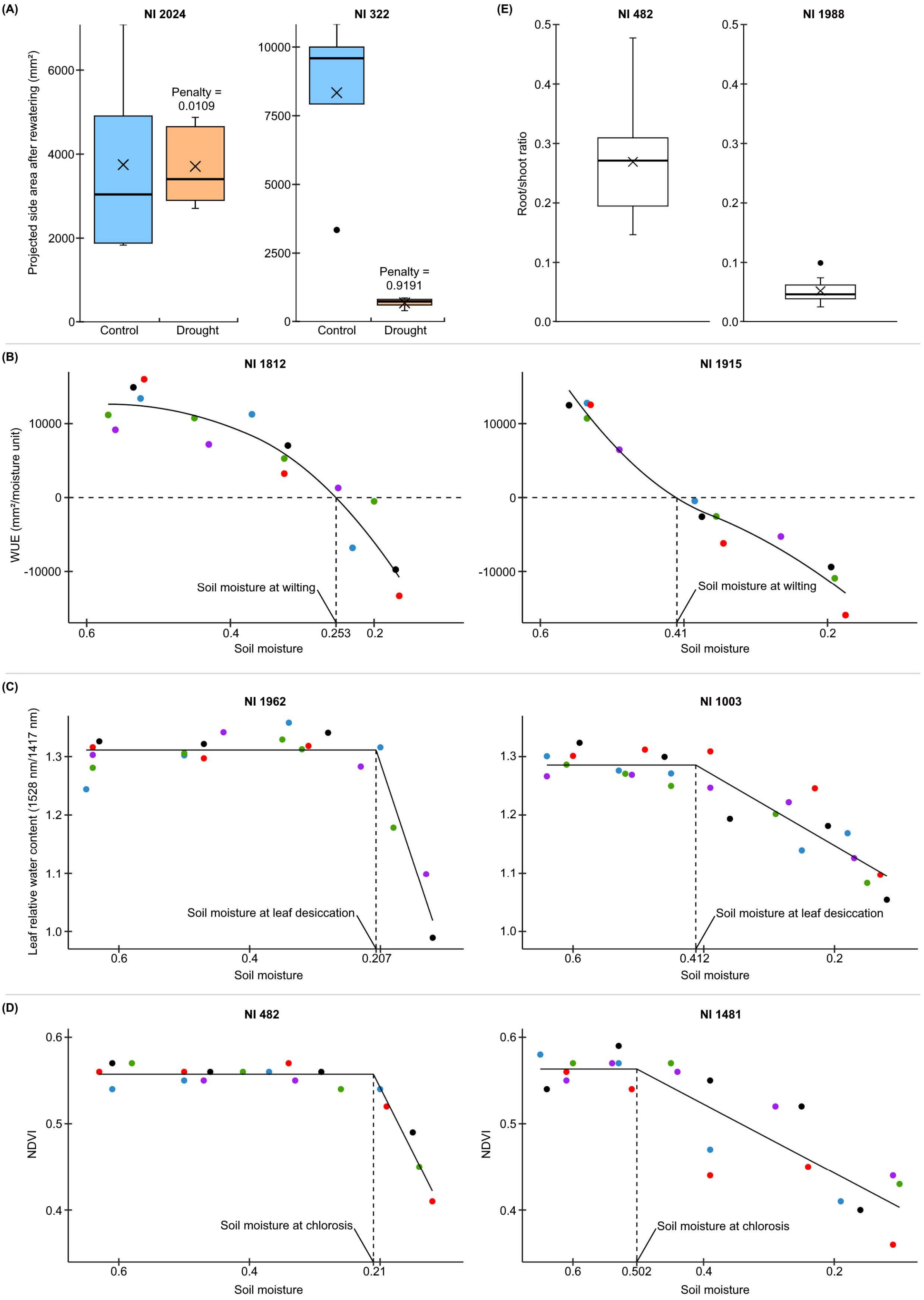

(A)
NI 2024
Projected side area after rewatering (mm²)
Penalty = 0.0109
Control
Drought
NI 322
Penalty = 0.9191
Control
Drought
(E)
NI 482
Root/shoot ratio
NI 1988
(B)
NI 1812
WUE (mm²/moisture unit)
Soil moisture at wilting
Soil moisture
NI 1915
Soil moisture at wilting
Soil moisture
(C)
NI 1962
Leaf relative water content (1528 nm/1417 nm)
Soil moisture at leaf desiccation
Soil moisture
NI 1003
Soil moisture at leaf desiccation
Soil moisture
(D)
NI 482
NDVI
Soil moisture at chlorosis
Soil moisture
NI 1481
NDVI
Soil moisture at chlorosis
Soil moisture

*Fig. S3.* Scatterplots of the five drought tolerance indicators in relation to each other.

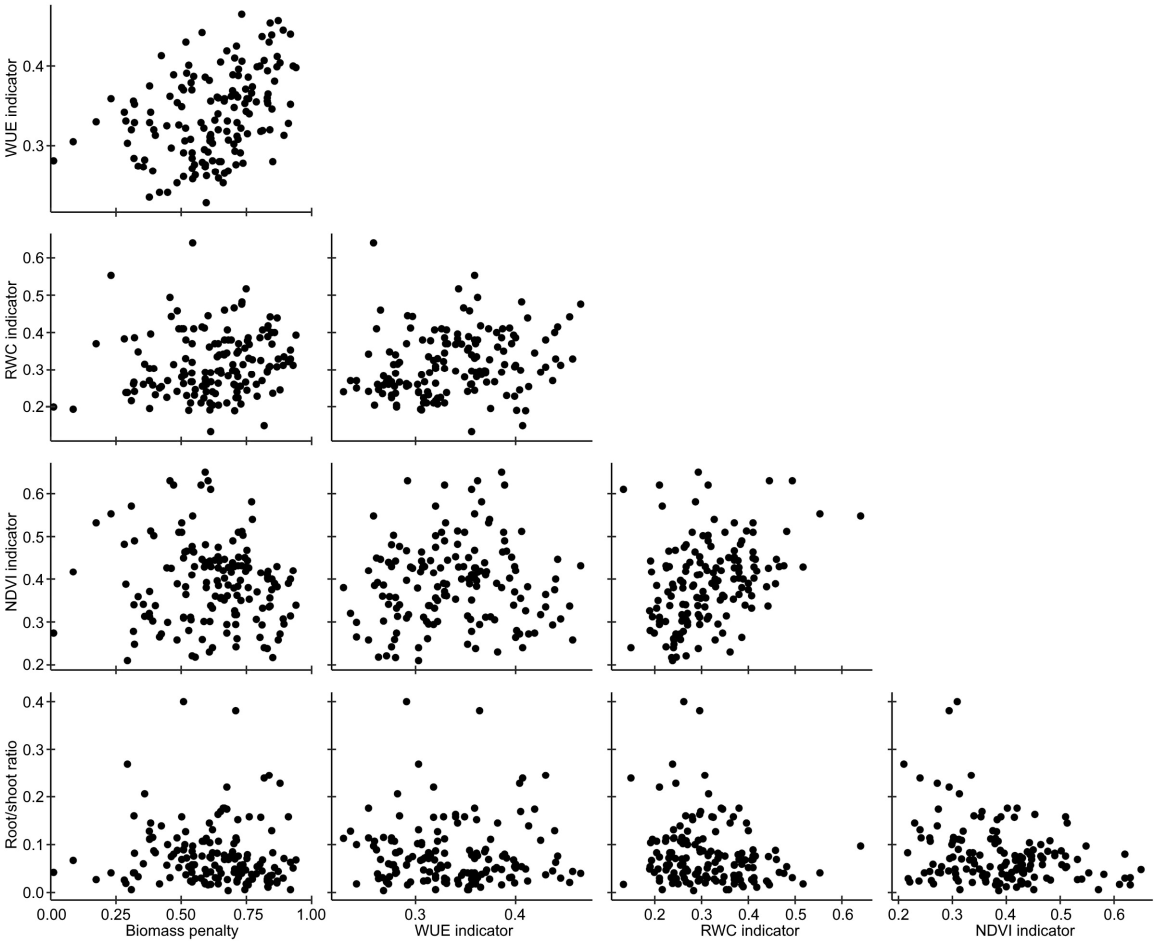

*Fig. S4.* Boxplots of median drought response indicator values for different subgenera of *Vigna* and *Phaseolus* beans (including former subgenera that were elevated to genera). Subgenera with 3 or fewer accessions were omitted. Black points indicate individual accessions. (A) Biomass penalty, (B) WUE indicator, (C) RWC indicator, (D) NDVI indicator, (E) Root/shoot ratio. N: number of accessions, SD: standard deviation, CV: coefficient of variation, *V.*: *Vigna*, *P.*: *Phaseolus*.

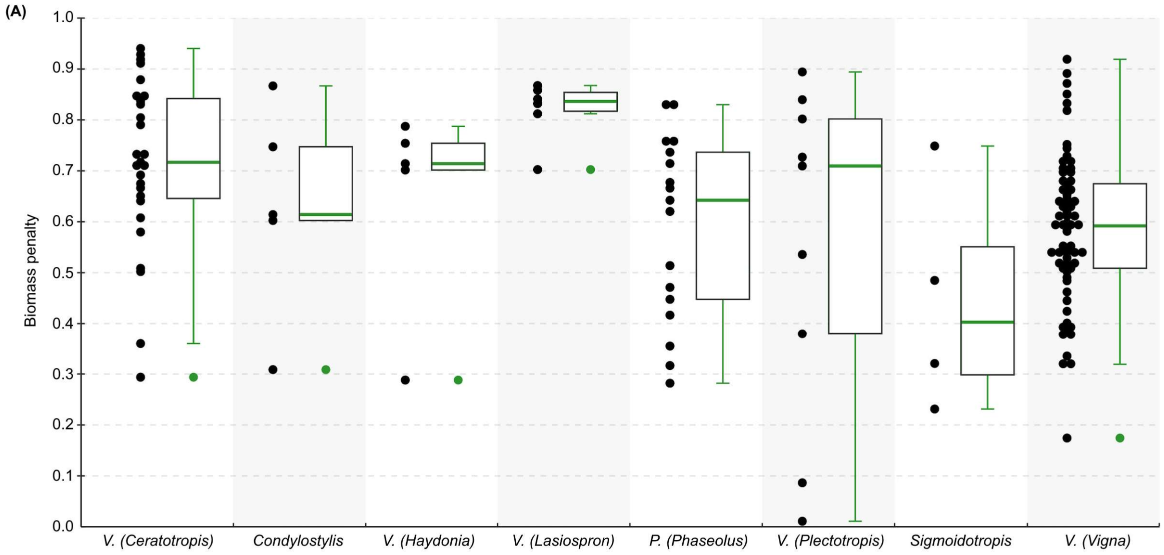


| | V. (Ceratotropis) | Condylostylis | V. (Haydonia) | V. (Lasiospron) | P. (Phaseolus) | V. (Plectotropis) | Sigmoidotropis | V. (Vigna) |
|---|---|---|---|---|---|---|---|---|
| N | 27 | 5 | 5 | 6 | 17 | 9 | 4 | 69 |
| Mean | 0.72 | 0.63 | 0.65 | 0.82 | 0.59 | 0.55 | 0.45 | 0.59 |
| SD | 0.17 | 0.21 | 0.20 | 0.06 | 0.18 | 0.33 | 0.23 | 0.15 |
| CV | 0.23 | 0.33 | 0.32 | 0.07 | 0.31 | 0.59 | 0.51 | 0.25 |

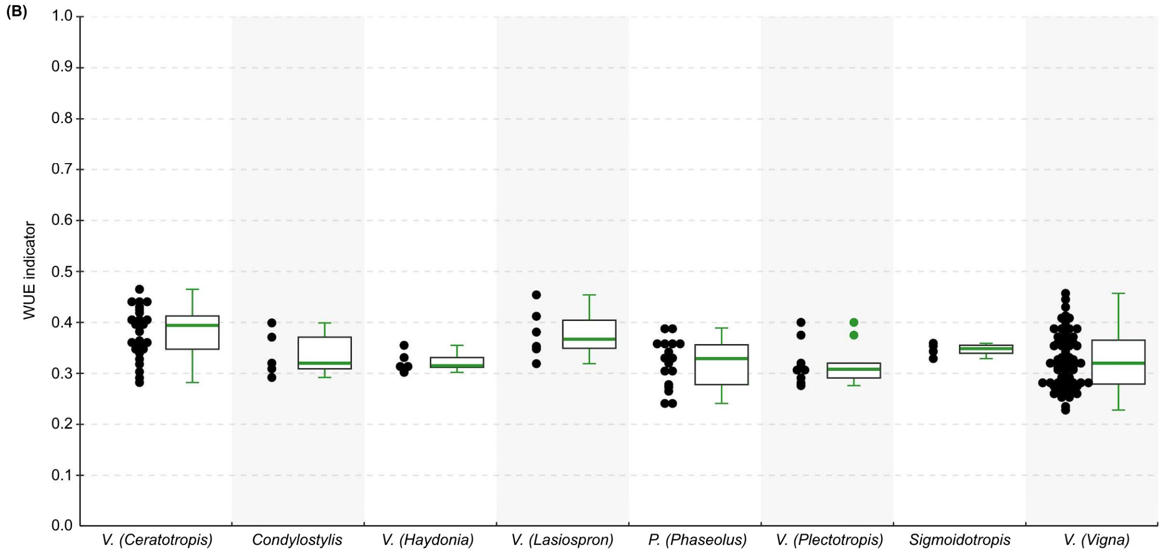


| | V. (Ceratotropis) | Condylostylis | V. (Haydonia) | V. (Lasiospron) | P. (Phaseolus) | V. (Plectotropis) | Sigmoidotropis | V. (Vigna) |
|---|---|---|---|---|---|---|---|---|
| N | 27 | 5 | 5 | 6 | 17 | 9 | 4 | 69 |
| Mean | 0.38 | 0.34 | 0.32 | 0.38 | 0.32 | 0.32 | 0.35 | 0.32 |
| SD | 0.05 | 0.04 | 0.02 | 0.05 | 0.05 | 0.04 | 0.01 | 0.05 |
| CV | 0.13 | 0.13 | 0.06 | 0.13 | 0.15 | 0.13 | 0.04 | 0.17 |

(C)

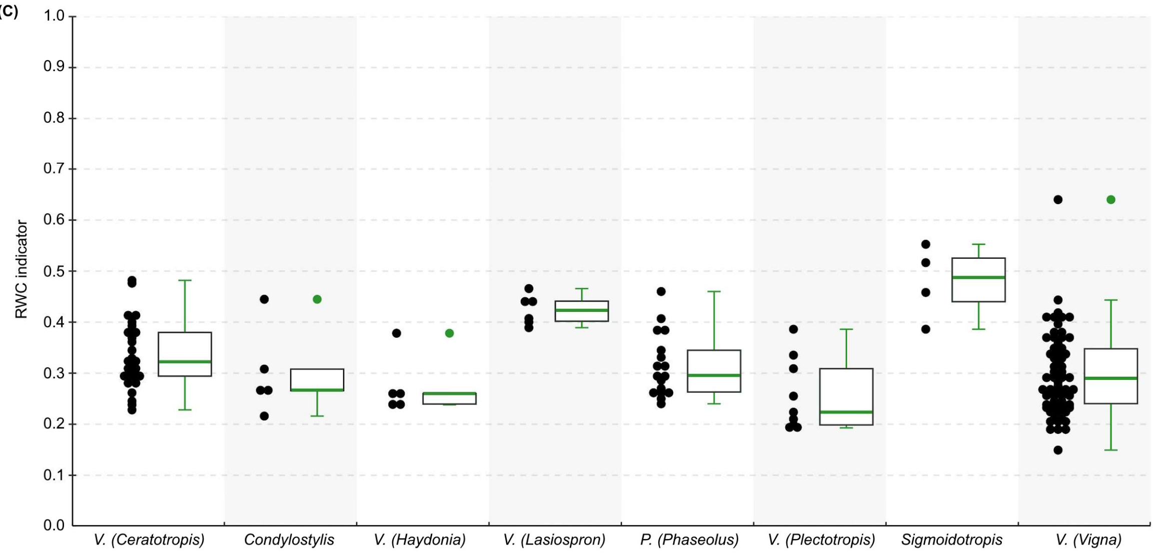


| | V. (Ceratotropis) | Condylostylis | V. (Haydonia) | V. (Lasiospron) | P. (Phaseolus) | V. (Plectotropis) | Sigmoidotropis | V. (Vigna) |
|---|---|---|---|---|---|---|---|---|
| N | 27 | 5 | 5 | 6 | 17 | 9 | 4 | 69 |
| Mean | 0.33 | 0.30 | 0.28 | 0.42 | 0.32 | 0.26 | 0.48 | 0.30 |
| SD | 0.07 | 0.09 | 0.06 | 0.03 | 0.06 | 0.07 | 0.07 | 0.08 |
| CV | 0.20 | 0.29 | 0.21 | 0.07 | 0.20 | 0.28 | 0.15 | 0.27 |

(D)

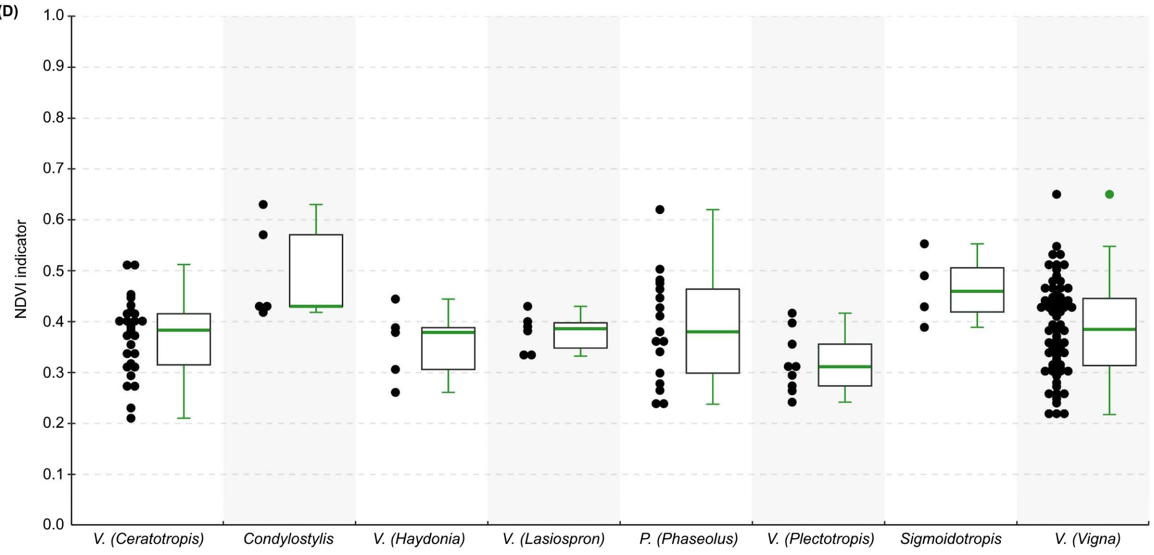


| | V. (Ceratotropis) | Condylostylis | V. (Haydonia) | V. (Lasiospron) | P. (Phaseolus) | V. (Plectotropis) | Sigmoidotropis | V. (Vigna) |
|---|---|---|---|---|---|---|---|---|
| N | 27 | 5 | 5 | 6 | 17 | 9 | 4 | 69 |
| Mean | 0.37 | 0.50 | 0.36 | 0.38 | 0.39 | 0.32 | 0.47 | 0.39 |
| SD | 0.08 | 0.10 | 0.07 | 0.04 | 0.11 | 0.06 | 0.07 | 0.09 |
| CV | 0.21 | 0.20 | 0.20 | 0.10 | 0.27 | 0.19 | 0.15 | 0.24 |

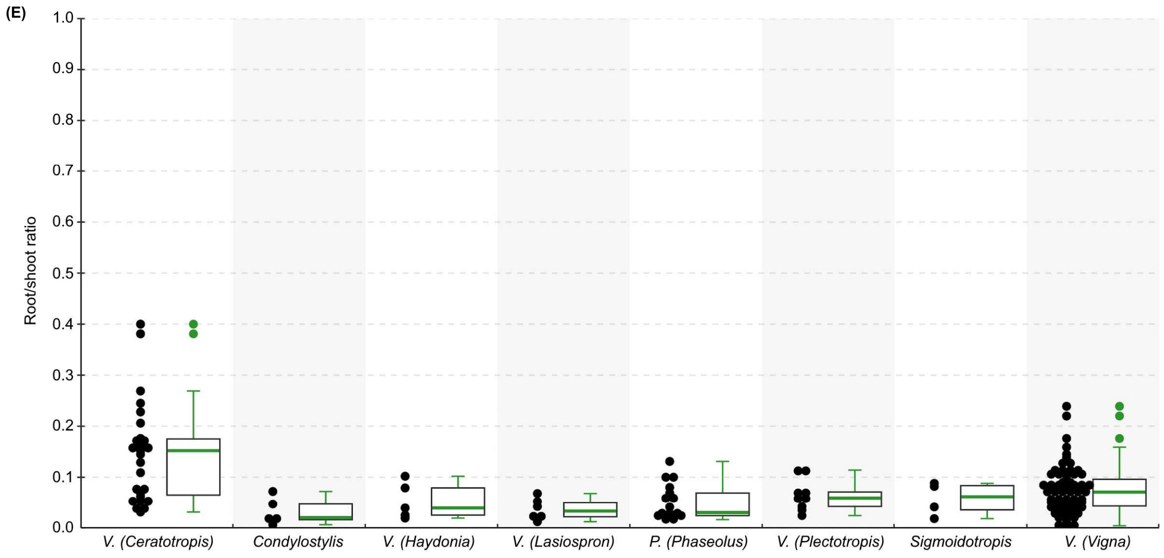


| | V. (Ceratotropis) | Condylostylis | V. (Haydonia) | V. (Lasiospron) | P. (Phaseolus) | V. (Plectotropis) | Sigmoidotropis | V. (Vigna) |
|---|---|---|---|---|---|---|---|---|
| N | 27 | 5 | 5 | 6 | 17 | 9 | 4 | 69 |
| Mean | 0.15 | 0.03 | 0.05 | 0.04 | 0.05 | 0.06 | 0.06 | 0.07 |
| SD | 0.10 | 0.03 | 0.04 | 0.02 | 0.03 | 0.03 | 0.03 | 0.05 |
| CV | 0.66 | 0.84 | 0.68 | 0.58 | 0.69 | 0.48 | 0.58 | 0.62 |

*Fig. S5*. Principal component analysis (PCA) performed with 151 accessions to visualise the relationships between five different drought response indicators. The distribution of the wild and cultivated accessions on the first two PC axes is indicated with red and blue dots, respectively.

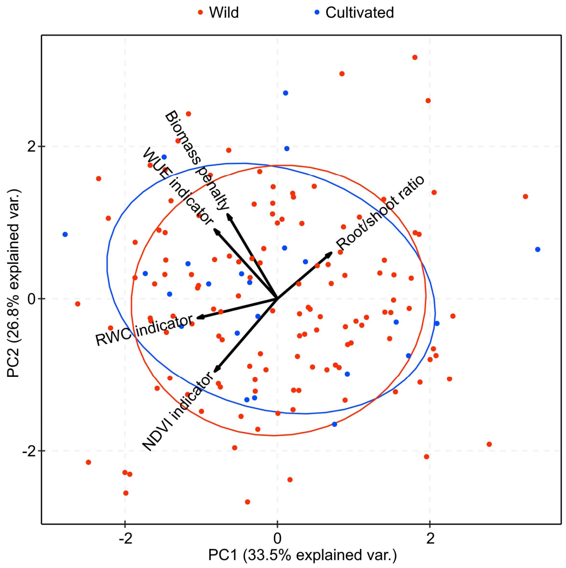
